# Intrinsic sensory disinhibition contributes to intrusive re-experiencing in combat veterans

**DOI:** 10.1101/602714

**Authors:** Kevin J. Clancy, Alejandro Albizu, Norman B. Schmidt, Wen Li

## Abstract

**ABSTRACT:** Intrusive re-experiencing of traumatic events is a hallmark symptom of posttraumatic stress disorder (PTSD). In contrast to abstract, verbal intrusions in other affective disorders, intrusive re-experiencing in PTSD is characterized by vivid sensory details as “flashbacks”. While prevailing PTSD models largely focus on dysregulated emotional processes, we hypothesize that deficient sensory inhibition in PTSD could drive overactivation of sensory representations of trauma memories, precipitating sensory-rich intrusions of trauma. In 86 combat veterans, we examined resting-state alpha (8-12 Hz) oscillatory activity (in both power and posterior→frontal connectivity), given its key role in sensory cortical inhibition, in association with intrusive re-experiencing symptoms. A subset (*N* = 35) of veterans further participated in an odor task (including both combat and non-combat odors) to assess olfactory trauma memory and emotional response. We observed a strong association between intrusive re-experiencing symptoms and attenuated resting-state posterior→frontal alpha connectivity, which were both correlated with olfactory trauma memory (but not emotional response). Importantly, olfactory trauma memory was further identified as a full mediator of the relationship between alpha connectivity and intrusive re-experiencing in these veterans, suggesting that deficits in intrinsic sensory inhibition can contribute to intrusive re-experiencing of trauma via heightened trauma memory. Therefore, by permitting unfiltered sensory cues to enter information processing and spontaneously activating sensory representations of trauma, impaired sensory inhibition can constitute a sensory mechanism of intrusive re-experiencing in PTSD.

*HIGHLIGHTS:*

- Alpha oscillations (indexing sensory inhibition) measured in 86 combat veterans
- Re-experiencing symptom severity was associated with attenuated alpha connectivity
- Trauma memory for, not emotional response to, odors mediated this relationship
- Trauma memories may arise via disinhibited activation of sensory representations
- Sensory systems may be novel target for intrusive re-experiencing symptom treatment

## 1. INTRODUCTION

Intrusive re-experiencing is a hallmark symptom of posttraumatic stress disorder (PTSD), characterized by the involuntary reactivation or reliving of trauma memories (American Psychiatric Association, 2013). To date, predominant models of PTSD have attributed these intrusive trauma memories to heightened emotion processing including excessive threat detection, exaggerated threat appraisal, and dysfunctional emotion regulation (Ehlers & Clark, 2000; Etkin & Wager, 2007; Liberzon & Abelson, 2016). Offering an alternative perspective, the dual representation theory of PTSD emphasizes dysregulated memory processes, implicating a sensory-bound representation system of threat memory (“S-memory”) in parallel to an abstract context-bound representation system (Brewin, 2014; Brewin, Gregory, Lipton, & Burgess, 2010). Furthermore, this S-memory system can be activated by basic sensory inputs, triggering the re-experiencing of trauma events in PTSD. This sensory-based memory system of trauma could account for re-experiencing symptoms in PTSD dominated by vivid, low-level sensory fragments of the trauma that are readily triggered by simple sensory cues (Ehlers, et al., 2002; Ehlers & Steil, 1995; Michael, Ehlers, & Halligan, 2005; Michael, Ehlers, Halligan, & Clark, 2005).

In keeping with this strong sensory feature in trauma re-experiencing, growing evidence indicates a sensory pathology in PTSD (Clancy, Ding, Bernat, Schmidt, & Li, 2017). Patients with PTSD often report being bothered or feeling ‘flooded’ by everyday stimuli, including background noises that others would not notice. Such complaints are further corroborated by systematic sensory profiling, which reveals sensory filtering/gating deficits and sensory hypersensitivity in these patients (Engel-Yeger, Palgy-Levin, & Lev-Wiesel, 2013; Stewart & White, 2008). Electrophysiological data also associate PTSD with poor repetition suppression to repeated auditory (“double-click”) stimuli (i.e., reduced P50 response reflecting impaired sensory gating) and exaggerated sensory evoked brain potentials as well as the mismatch negativity (reflecting sensory cortical hyperactivity) in response to simple, neutral stimuli (Javanbakht, Liberzon, Amirsadri, Gjini, & Boutros, 2011; Morgan & Grillon, 1999; Neylan, et al., 1999; Stewart & White, 2008). Therefore, it is plausible that deficient sensory gating and heightened sensory activity in PTSD could over-activate the sensory-based trauma memory system by permitting excessive, unfiltered sensory inputs into information processing and turning on stored sensory representations of trauma.

A key neural mechanism underlying sensory gating and sensory cortical regulation involves alpha-frequency (8-12 Hz) oscillations, which dominate neural synchrony in the awake restful state (Bazanova & Vernon, 2014; Buzsaki, Logothetis, & Singer, 2013; Klimesch, 2012; Palva & Palva, 2007). Specifically, alpha oscillatory activity exerts inhibitory influences on sensory cortical neuronal firing and excitation (Foxe & Snyder, 2011; Jensen & Bonnefond, 2013; Klimesch, Sauseng, & Hanslmayr, 2007). Furthermore, via long-range, posterior→frontal projections, alpha oscillations mediate inhibitory bottom-up information flow from the sensory cortex to frontal regions to influence global neural activity (Hillebrand, et al., 2016; Johnson, et al., 2017; Sadaghiani & Kleinschmidt, 2016; Tang, et al., 2007). This long-range alpha connectivity plays a critical role in gating the entry of sensory input into downstream processing and, eventually, conscious awareness, thereby regulating perception, imagery, and working memory (Doesburg, Green, McDonald, & Ward, 2009; Jensen, Bonnefond, & VanRullen, 2012; Mathewson, et al., 2011; Samaha, Boutonnet, Postle, & Lupyan, 2018; Samaha & Postle, 2015). Alternatively, deficient alpha activity could compromise this restricted access such that unfiltered sensory input would flood information processing, activating stored memory representations and eliciting vivid imagery and perception, akin to intrusive re-experiencing in PTSD. In fact, we have recently demonstrated severely suppressed alpha activity in patients with PTSD, including both local alpha power and posterior→frontal alpha connectivity (Clancy, et al., 2017). Notably, deficient posterior→frontal alpha connectivity was directly associated with excessive frontal cortical activity and disinhibition symptoms. As such, we hypothesized that deficient alpha activity in PTSD could serve as a key mechanism for intrusive re-experiencing by impeding sensory inhibition and thereby over-activating sensory representations of trauma memory.

To test this hypothesis, we recruited a sample of combat-exposed veterans (*N* = 86) and examined their intrusive re-experiencing symptoms in relation to alpha activity, including alpha power and alpha posterior→frontal connectivity (using Granger causality analysis) (Ding, Chen, & Bressler, 2006). Akin to the spontaneous, involuntary nature of intrusions, we focused on intrinsic alpha activity as measured during the resting-state. In a subsample of the veterans (*N* = 35), we further tapped into sensory-based activation of autobiographical memory of combat experience. We chose olfactory (combat and non-combat) cues to activate olfactory trauma memory given that odors are known to activate vivid, sensory-rich memories in a Proustian manner (Herz & Schooler, 2002; Wilson & Stevenson, 2003) and strongly trigger trauma memories in PTSD (Cortese, Leslie, & Uhde, 2015; Vermetten & Bremner, 2003; Vermetten, Bremner, Skelton, & Spiegel, 2007). As illustrated in Figure 1, we tested our hypothesis in a mediation model, where intrinsic alpha deficits contributed to intrusive re-experiencing through overactivation of sensory-based trauma memory. To disambiguate this memory mechanism from the potentially competing mechanism of exaggerated threat processing (as prominently implicated in PTSD) (Liberzon & Abelson, 2016), we also assessed emotional response to odors to pit that against olfactory trauma memory in relation to alpha attenuation and intrusive re-experiencing.

**Figure 1.**
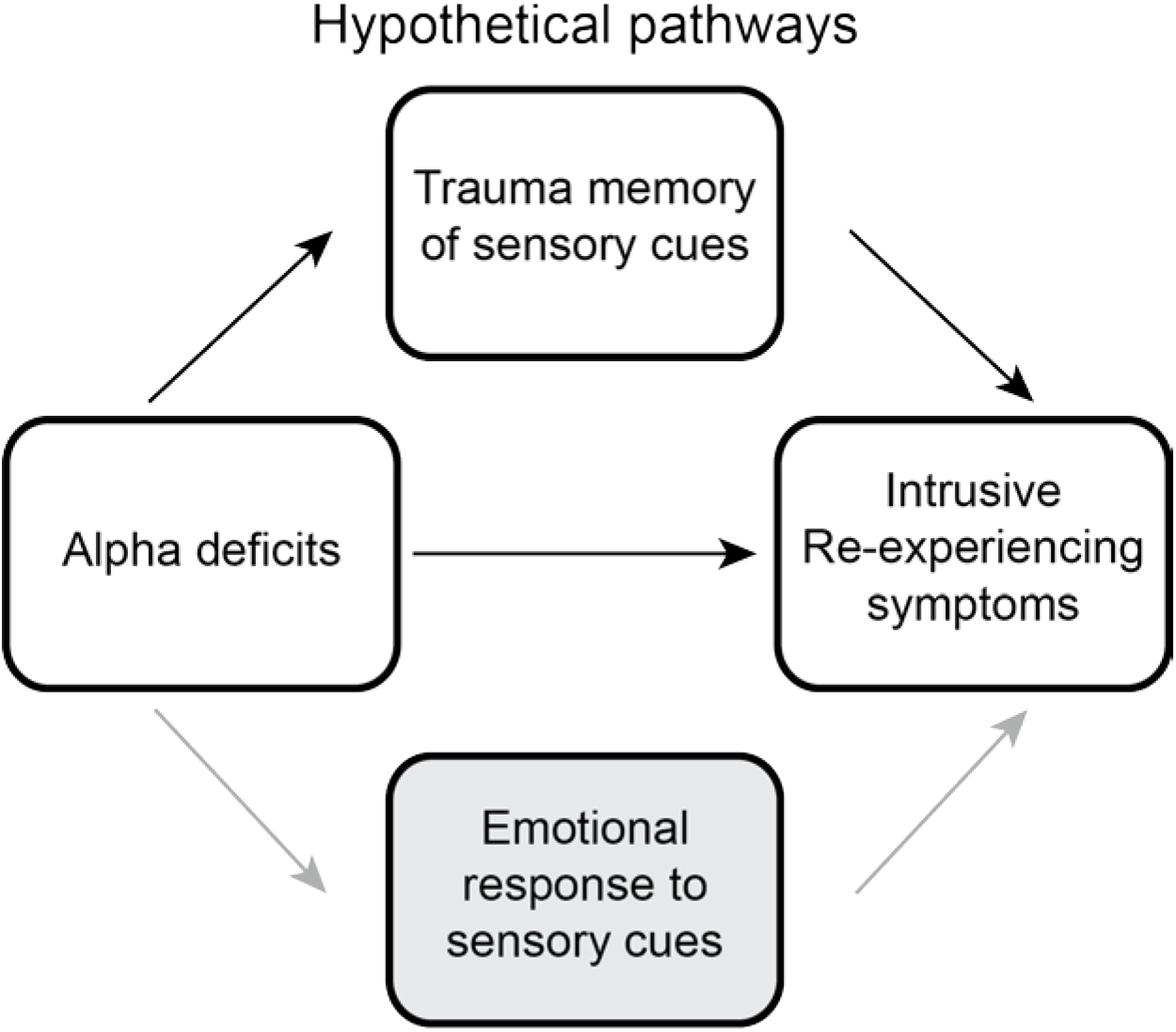
A sensory model for intrusive re-experiencing symptoms, testing the hypothesis that deficient inhibition of sensory information increases intrusive re-experiencing symptoms by increasing trauma memory recall. Emotional response to sensory cues was considered as a competing mediator in this model.

## 2. METHODS

### 2.1. Participants

Ninety-two combat-exposed veterans participated in the study after providing informed consent approved by the Florida State University Institutional Review Board and the Department of Defense Human Research Protection Official’s Review. Participants had no history of severe neurological disorders or traumatic brain injury, no current or past psychotic spectrum or bipolar disorders, and no current substance dependence or abuse of opioids, stimulants, or cocaine. Six participants were excluded due to excessive EEG artefact or failure to follow instructions, resulting in a final sample of eighty-six veterans (mean age: 45.9 ± 12.6 years; 10 females; see Table 1 for demographics).

**Table 1:**
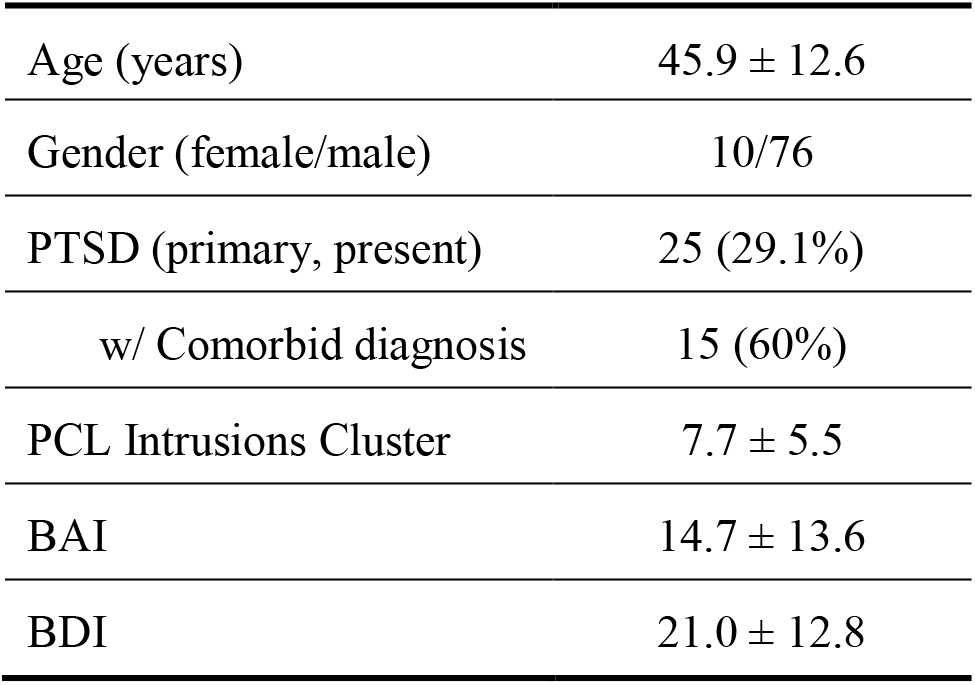
Participant Demographics (*n* = 86). PCL = Posttraumatic Stress Disorder Checklist; BAI = Beck Anxiety Inventory; BDI = Beck Depression Inventory

### 2.2. Clinical Assessment

All participants underwent the PTSD module of the Structured Clinical Interview for DSM-V (American Psychiatric Association, 2013), among which 25 met the DSM-V criteria for current PTSD. Participants also completed the PTSD Checklist (PCL) (Blevins, Weathers, Davis, Witte, & Domino, 2015), from which we summed the responses to Items 1-5 that assess symptoms of intrusions to generate a total score of intrusive re-experiencing symptom severity (range: 0-20). Additionally, participants completed the Beck Depression Inventory (BDI-II) (Beck, Steer, & Brown, 1996) and Beck Anxiety Inventory (BAI) (Beck, Epstein, Brown, & Steer, 1988) to index general depression and anxiety symptoms. Scores on these measures are provided in Table 1.

### 2.3. EEG acquisition and analyses

EEG data were recorded from a 96-channel BrainProducts actiChamp system (1000 Hz sampling rate, 0.05 – 200 Hz online bandpass filter, referenced to the FCz channel) for 2 minutes, with eyes fixated on the central crosshair of a computer screen. Electro-oculogram (EOG) was recorded using four electrodes with vertical and horizontal bipolar derivations. EEG/EOG data were downsampled to 250 Hz, high-pass (1 Hz) and notch (60 Hz) filtered, and re-referenced to the average of all EEG channels. We applied the *Fully Automated Statistical Thresholding for EEG artifact Rejection* (FASTER) algorithm for artifact detection and rejection (Nolan, Whelan, & Reilly, 2010).

#### 2.3.1. Power analyses

EEG oscillation power was computed for individual channels for each epoch (1-sec) using the multitaper spectral estimation technique (Mitra & Pesaran, 1999). Alpha (8-12 Hz) power was normalized by the mean power for the global spectrum (1-50 Hz) within each epoch, and averaged across occipito-parietal electrodes, where alpha is maximally distributed (Foxe & Snyder, 2011; Klimesch, 2012; Palva & Palva, 2007) (Figure 2A). Figure 2A further illustrates the intracranial alpha source, localized to the striate/extrastriate visual cortices (e.g., maximum in the cuneus; x, y, z = −10, −80, 10) based on the exact Low-Resolution Electromagnetic Tomography (eLORETA) (Pascual-Marqui, et al., 2011), in keeping with the literature (Cuspineda, et al., 2009; Neuner, et al., 2014; Pascual-Marqui, et al., 2011).

**Figure 2.**
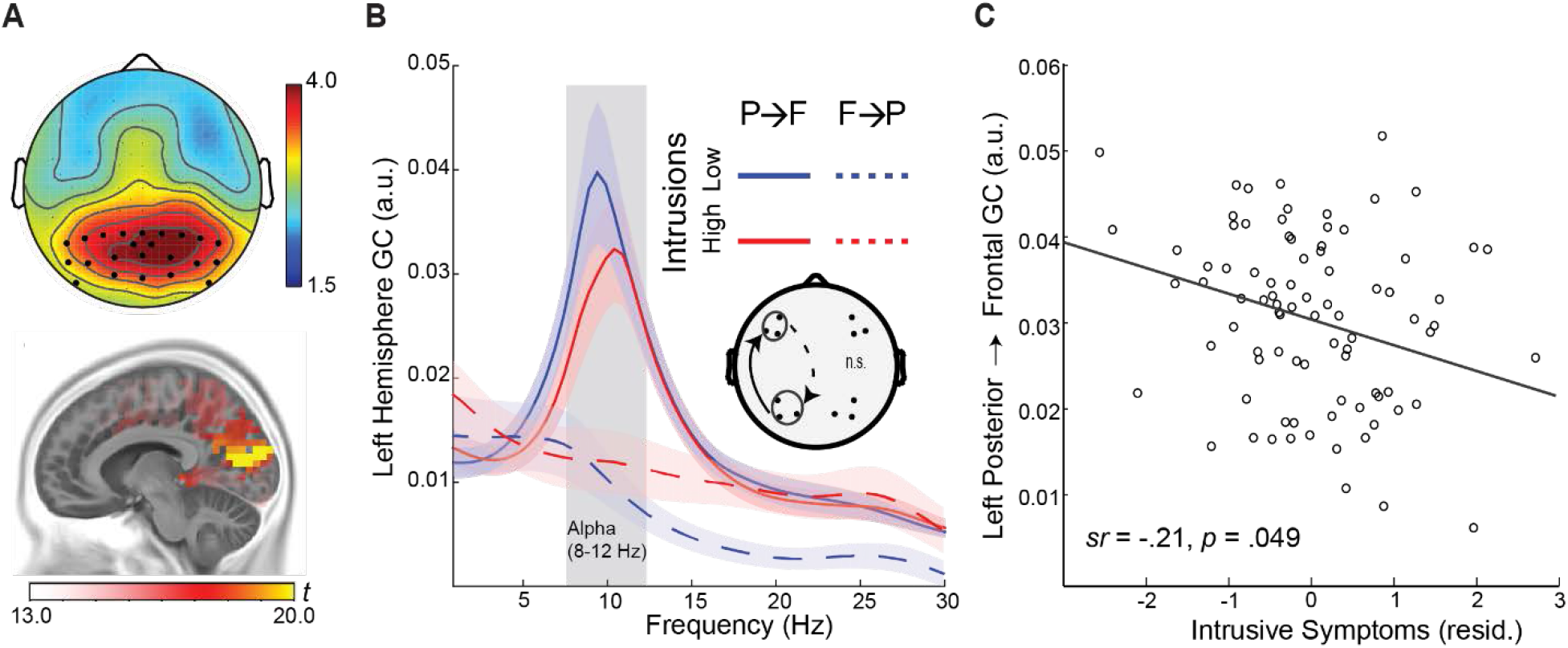
Attenuated left-hemisphere alpha connectivity was related to greater intrusion symptoms. A) Topography of alpha power, maximally distributed over occipitoparietal electrodes. Intracranial sources of alpha power were localized to the striate/extrastriate visual cortices (e.g., maximum in the cuneus; x, y, z = −10, −80, 10). B) Spectra of left-hemisphere Granger causality for participants with high vs. low intrusive re-experiencing symptoms (median split), demonstrating reduced alpha (8-12 Hz) posterior→frontal connectivity in high-intrusion participants. The opposite, frontal→posterior GC was minimal and equivalent for the two groups. Shaded ribbons = standard error of the mean (SEM). C) Negative correlation between alpha connectivity and intrusive re-experiencing symptoms, controlling for depression symptoms (BDI scores). P→F = posterior→frontal; F→P = frontal→posterior.

#### 2.3.2. Directed alphafrequency connectivity (Granger causality) analyses

Alpha-frequency Granger causality (GC) analysis (Ding, et al., 2006; Geweke, 1982) was performed to assess posterior→frontal causal connectivity in the alpha band. Following transformation into reference-free, current source density (CSD) data using the surface Laplacian algorithm (Nunez, et al., 1997; Perrin, Pernier, Bertrand, & Echallier, 1989; Wang, Rajagovindan, Han, & Ding, 2016), EEG data from ipsilateral posterior-frontal pairs were submitted to bivariate autoregressive (AR) modeling, from which Granger causality spectra were derived (Ding, Bressler, Yang, & Liang, 2000; Ding, et al., 2006). A model order of 20 (80 ms in time for a sampling rate of 250 Hz) was chosen in a two-step process: (1) Akaike Information Criterion (AIC) and (2) comparing the spectral estimates obtained by the AR model and that by the Fourier based method for data pooled across all subjects (Wang, et al., 2016). Ipsilateral posterior-frontal electrode pairs were selected *a priori* based on previous studies (Clancy, et al., 2017; Johnson, et al., 2017). Akin to the dominance of the posterior→frontal directed propagation of alpha oscillations at rest (Engel, Fries, & Singer, 2001; Hillebrand, et al., 2016; Tang, et al., 2007), minimal frontal→posterior alpha connectivity was observed (Fig 2B).

### 2.4. Odor task

A subsample of thirty-five veterans (mean age: 42.6 ± 13.4 years; 4 females) then participated in an odor task, whereby six combat and six non-combat odors contained in amber glass bottles were presented. Combat odors included two variations of spent gunpowder and burning rubber, as well as the smell of burning flesh and of body decay (ScentAir^™^, NC). Non-combat odors included neutral (or mildly pleasant) chemicals: cleaning fluid (ScentAir^™^, NC), acetophenone, eugenol, cedrol, alpha-ionone, and citronellol (Fisher Scientific, NH). Both combat and non-combat odors were rated to be of equal, moderate intensity [mean (SD) = combat: 52.2 (12.7) versus non-combat: 48.7 (11.8); *t*_1,34_ = 1.65, *p* = .11].

To assess the level of trauma memory activation by sensory cues, we measured the degree to which the odors were recognized as being experienced during the veterans’ combat deployment. To disambiguate memory and emotion processes, we also assessed the level of emotional responses elicited by the odors. Specifically, on each odor presentation (randomized across the 12 odors), veteran participants were asked to rate on a visual analog scale (VAS; 0-100) how strongly the odor was associated with and how frequently it was experienced during their combat deployment. These two ratings were averaged into an olfactory trauma memory score. Participants were also asked to rate how strongly the odor elicited distress, disgust, and anxious arousal, which were averaged into an olfactory emotional response score. The order of these ratings was randomized across odors. Paired samples t-tests revealed that combat odors, relative to non-combat odors, had higher trauma memory scores [mean (SD) = combat odors: 32.83 (14.81) versus non-combat odors: 25.31 (13.85); *t*_1,34_ = 4.21, *p* < .001], and higher emotional response scores [combat odors: 43.46 (15.76) versus non-combat odors: 30.04 (14.99); *t*_1, 34_ = 7.36, *p* < .001], confirming their distinct odor categories. Finally, trauma memory and emotional response scores were marginally correlated (combat odor: *r* = .32, *p* = .063; non-combat odor: *r* = .33, *p* = .051], indicating that they reflected related but distinct constructs.

### 2.5. Statistical Analyses

Multiple regression analyses were performed to examine associations between alpha activity (power and GC) and intrusive re-experiencing symptoms in the general sample, with BDI scores entered as covariates to isolate PTSD symptoms from comorbid depressive symptoms. Significant correlations between alpha activity (power or GC) and intrusive symptoms in the general sample would then motivate omnibus repeated measures analyses of covariance (rANCOVAs) in the subsample to examine the relation between these variables and olfactory trauma memory and emotional response. Specifically, omnibus rANCOVAs were performed on the olfactory scores with Category (combat vs. non-combat) and Response (combat memory vs. emotional response) as independent variables and alpha activity (power or GC value) as a covariate. “Response” was entered as an independent variable in the ANCOVA to pit olfactory trauma memory against emotional response, thereby isolating specific memory effects. In addition, a similar ANCOVA was conducted with the severity of intrusive re-experiencing symptoms (based on the PCL) as a covariate to examine its association with olfactory trauma memory and emotional response. Provided that these variables were correlated with each other, mediation analyses were conducted to test the hypothesis of trauma memory or emotional response mediating the relationship between alpha activity and intrusive re-experiencing symptoms. The PROCESS macro for SPSS (Hayes, 2012) was used to estimate 5,000 bias-corrected bootstrap samples, from which a 95% confidence interval (CI) was created to test the indirect effect of alpha activity on re-experiencing symptoms through olfactory processes. Again, depression symptoms (BDI scores) were entered as covariates to control for effects of comorbid depression.

## 3. RESULTS

### 3.1. Alpha power

A multiple regression analysis revealed no association between posterior alpha power and intrusive symptoms, after controlling for depression symptoms (*p* = .640). Given this null effect, the follow-up rANCOVAs were not justified and thus not reported here. For reference, we presented results of the rANCOVAs with alpha power in the supplemental material.

### 3.2. Alpha posterior→frontal causal connectivity

#### 3.2.1. Attenuated alpha connectivity associated with greater intrusive re-experiencing

Multiple regression analyses revealed a negative association between alpha GC and intrusive re-experiencing, significantly in the left hemisphere (*sr* = −.21, *p* = .049) and weakly in the right hemisphere (*sr* = −.01, *p* = .371), after controlling for depression symptoms (Figure 2). This association persisted in the subsample of 35 veterans that completed the odor task (left hemisphere: *sr* = −.43, *p* = .010; right hemisphere: *sr* = −.09, *p* = .629). Therefore, veterans with attenuated left-hemispheric alpha connectivity showed greater intrusive re-experiencing symptoms. With this association between alpha connectivity and intrusion symptoms, we conducted the following ANCOVAs (limited to the left hemisphere that exhibited significant effects).

#### 3.2.2. Olfactory trauma memory associated with greater intrusive re-experiencing symptoms

An omnibus rANCOVA of Category (combat vs. non-combat) X Response (memory vs. emotion) X intrusive re-experiencing symptoms on olfactory scores revealed a trending 3-way interaction of Category X Response X intrusive-re-experiencing (*F*_1, 33_ = 3.25, *p* = .080, *ŋ*_p_^2^ = .09). Breaking down the interaction by Response, a follow-up rANCOVA of Category X intrusive re-experiencing for olfactory emotional response scores yielded a trending effect of Category X intrusive re-experiencing (*F*_1,33_ = 2.88, *p* = .099, *ŋ*_p_^2^ = .08): intrusive symptoms were correlated with emotional response to combat odors (*sr* = .35, *p* = .041) but not with non-combat odors (*sr* = .16, *p* = .355; Figure 3A). A similar rANCOVA of Category X intrusive re-experiencing for olfactory trauma memory scores showed a main effect of intrusive re-experiencing (*F*_1, 33_ = 11.22, *p* = .002, *ŋ*_p_^2^ = .25), but no Category X intrusive re-experiencing interaction (*p* = .529). That is, as illustrated in Figure 3B, trauma memory for both combat (*sr* = .41, *p* = .013) and non-combat (*sr* = .53, *p* = .001) odors was significantly associated with intrusive symptoms. Therefore, individuals with greater trauma memory (for both combat and non-combat cues) exhibited greater intrusive symptoms.

**Figure 3.**
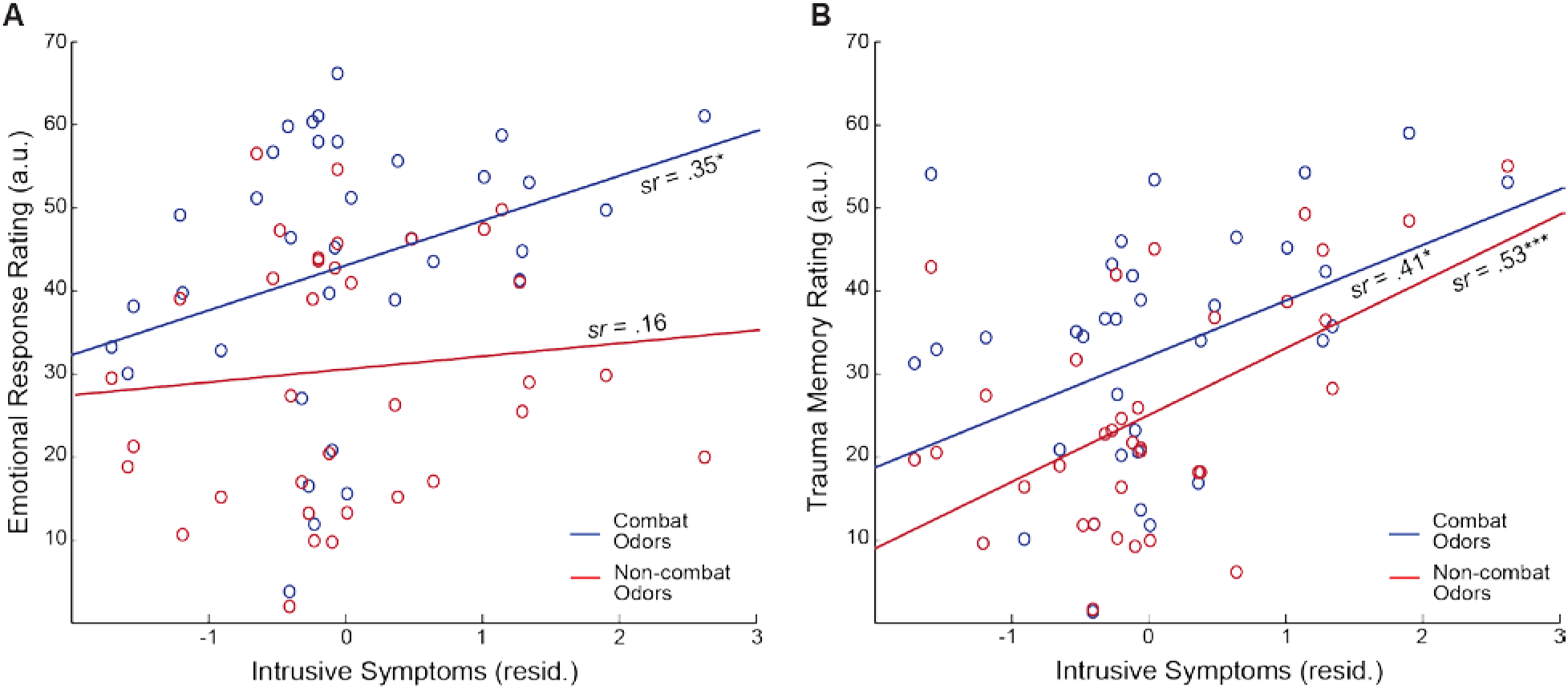
Olfactory responses were associated with intrusion symptoms. A) Greater emotional response to combat, but not non-combat, odors was associated with greater intrusive re-experiencing symptoms (controlling for depression). B) Greater trauma memory for both combat and non-combat odors was associated with greater intrusive re-experiencing symptoms. \**p* < .05; \*\**p* < .01; \*\*\**p* < .005; † *p* < .1.

#### 3.2.3. Olfactory trauma memory associated w ith decreased alpha connectivity

To examine the effect of alpha connectivity, a similar omnibus rANCOVA (Category x Response x alpha GC) on olfactory scores revealed a significant interaction between Response and alpha GC (*F*_1,33_ = 5.19, *p = .029, *ŋ*_p_^2^* = .14): reduced alpha connectivity was associated with greater olfactory trauma memory (*sr* = −.40, *p* = .018; Figure 4A) but not olfactory emotional response (*p* = .839; Figure 4B). There was no Category effect or Category-by-Response interaction (*p*’s > .929), suggesting that the association of trauma memory with alpha connectivity spanned across combat (*sr* = −.36, *p* = .035) and non-combat (*sr* = −.39, *p* = .023) odors (Figure 4A). Therefore, veterans with reduced alpha posterior→frontal connectivity also exhibited heightened olfactory trauma memory (but not emotional response).

**Figure 4.**
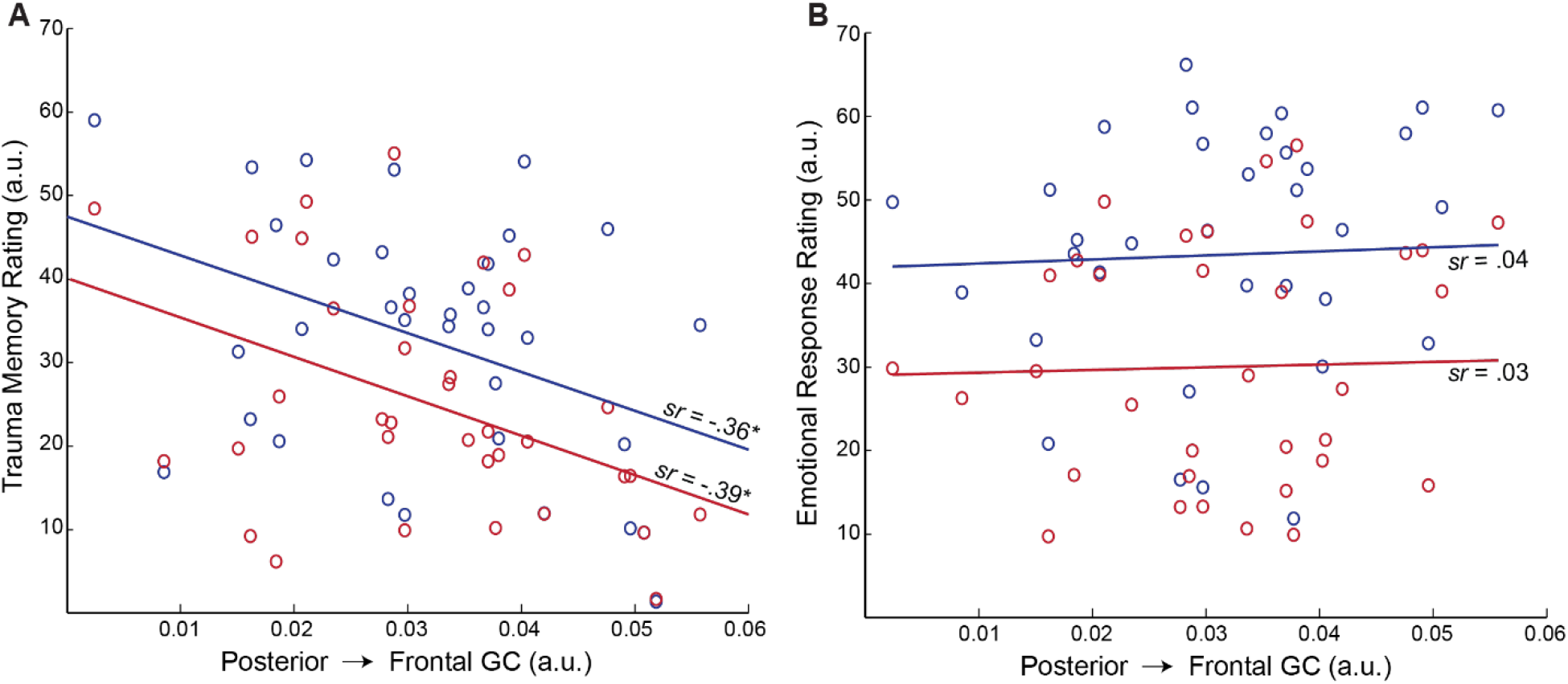
Attenuated alpha connectivity was related to heightened trauma memory for olfactory cues. A) Weakened left-hemisphere posterior→frontal Granger causality was associated with greater trauma memory for both combat and non-combat odors. B) Left-hemisphere posterior→frontal Granger causality demonstrated no association with emotional response to neither combat nor non-combat odors. \**p* < .05.

#### 3.2.4. Olfactory trauma memory mediating the association between alpha connectivity and intrusive re-experiencing

To further assess the memory mechanism through which attenuated alpha activity was related to greater intrusive re-experiencing symptoms, a mediation analysis was performed between alpha posterior→frontal GC and intrusive re-experiencing symptoms, with olfactory trauma memory (combat and non-combat collapsed) serving as the mediator. Bias-corrected bootstrap mediation analysis revealed a full mediation by olfactory trauma memory (indirect effect: *β* = −.16, SE = .11, 95% CI = [−.500, −.011]), such that weakened alpha connectivity increased re-experiencing symptoms through heightened trauma memory for both non-combat and combat cues (Figure 5).

**Figure 5.**
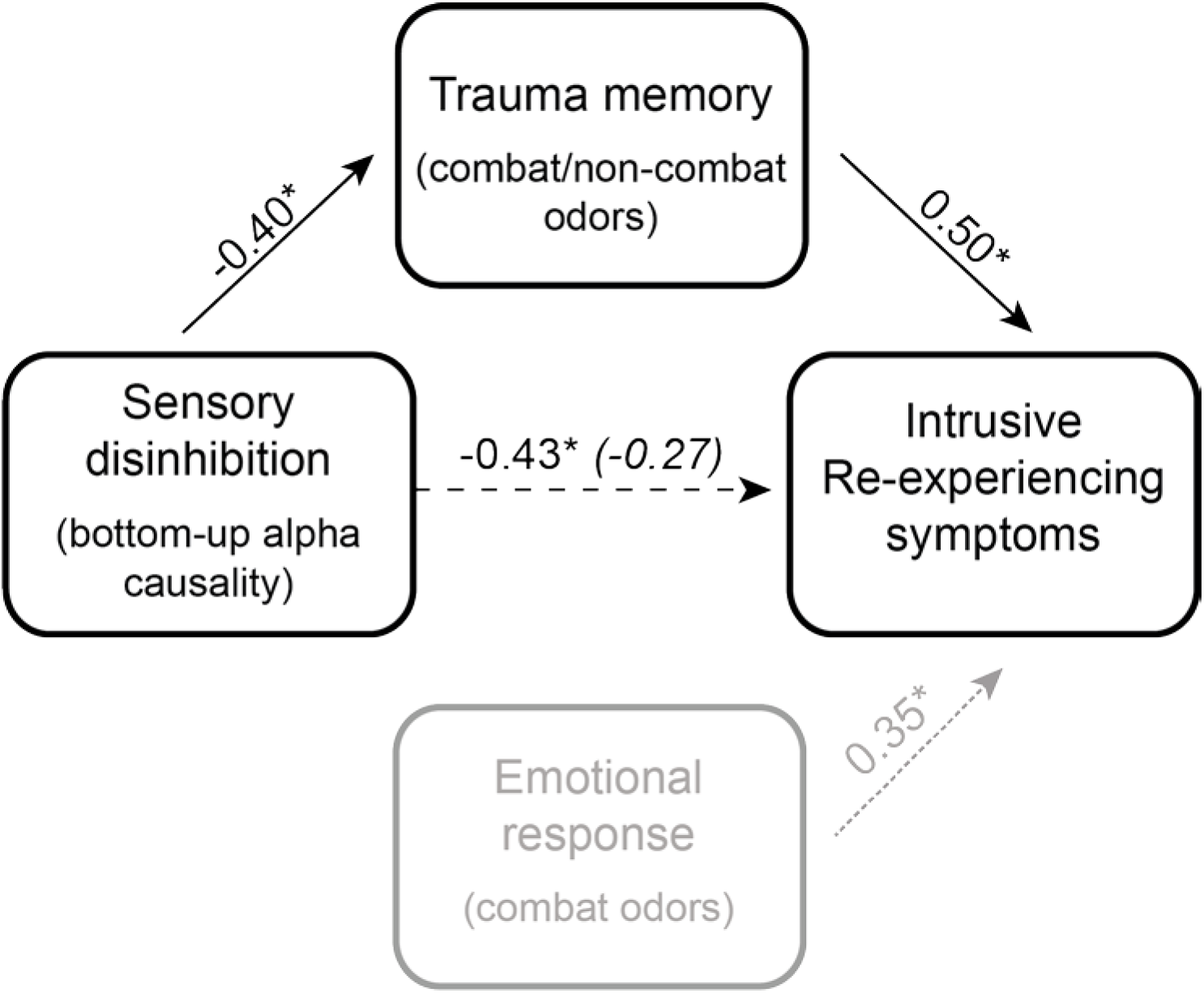
Mediation: Trauma memory for olfactory cues fully mediated the relationship between alpha connectivity and intrusive re-experiencing symptoms, such that the correlation decreased significant and became nonsignificant following the inclusion of trauma memory scores. This memory-related mediation contrasts to emotional response, which was not associated with alpha activity (and so not included in the mediation analysis). The path strengths are indicated by beta coefficients. The parenthetical beta coefficient reflects the direct path strength after controlling for trauma memory. \**p* < .05.

## 4. DISCUSSION

In a sample of combat veterans, we demonstrated an association between heightened intrusive re-experiencing symptoms and attenuated resting-state alpha posterior→frontal connectivity. We also observed that exaggerated olfactory trauma memory not only correlate with intrusive symptom severity and alpha connectivity (beyond olfactory emotional response), it also mediated their association. Intrusive re-experiencing in PTSD reflects the involuntary recall or re-living of traumatic events characterized by vivid, sensory-rich details (Ehlers, et al., 2002; Michael, Ehlers, & Halligan, 2005; Michael, Ehlers, Halligan, et al., 2005), while attenuated resting-state alpha posterior→frontal connectivity reflects intrinsic disinhibition of sensory propagation to higher-order regions. Therefore, these results suggest that intrinsic sensory disinhibition can contribute to intrusive re-experiencing symptoms through the overactivation of sensory trauma memory. Taken together, the current findings highlight a specific sensory mechanism underlying symptoms of intrusive trauma re-experiencing, lending credence to a sensory pathology of PTSD.

Dominant models of PTSD have largely focused on excessive threat response (e.g., detection and appraisal) and deficient emotion regulation, whereby traumatic memory and intrusive re-experiencing arise via heightened threat processing and attenuated executive control (Ehlers & Clark, 2000; Etkin & Wager, 2007; Liberzon & Abelson, 2016; Rauch, Shin, & Phelps, 2006). Indeed, we observed a significant relation between exaggerated emotional response to combat odors and intrusive symptoms. However, pitting emotional response against trauma memory for the odors (as a factor of Response in the ANOVAs reported above), we observed that intrusive re-experiencing symptoms were particularly associated with exaggerated trauma memory. This selective association was further corroborated by a multiple regression analysis with both emotional response and trauma memory as regressors, indicating a unique relation between trauma memory and intrusive symptoms (*sr* = .45, *p* < .01) over and beyond emotional response (*sr* = .12, *p* = .50; see Supplemental Materials). That is, beyond a general emotional bias in PTSD, exaggerated trauma memory makes a unique contribution to intrusive symptoms.

This exaggerated trauma memory in veterans with severe intrusive symptoms extended to both combat and non-combat odors. Strikingly, trauma memory for non-combat odors was predictive of greater intrusive re-experiencing symptoms over and beyond trauma memory for combat odors (*sr* = .36, *p* = .04; *sr* = .05, *p* = .78, respectively; see Supplemental Materials). This false recognition for non-combat odors, which were neutral/mildly pleasant, concurs with the commonly observed false memory of non-trauma-related cues in individuals with PTSD (Brewin, Huntley, & Whalley, 2012; Geraerts, et al., 2009) and clinical manifestations of intrusive memories triggered by a wide range of cues, including those bearing little association with the trauma (Ehlers, 2010). This pattern of effects also aligns with the dual representation theory of PTSD such that simple sensory cues alone, independent of context, would suffice to trigger involuntary information processing and thereby activate sensory representations of trauma memory and trigger intrusive re-experiencing (Brewin, 2014; Brewin, et al., 2010).

Furthermore, these findings coincide with the growing evidence of sensory disinhibition in PTSD in that the sensory system fails to filter and block irrelevant or non-salient (e.g., non-trauma) sensory cues from entering information processing (Clancy, et al., 2017; Neylan, et al., 1999; Stewart & White, 2008). By suppressing neural excitability and modulating long-range cortico-cortical communication, alpha oscillations serve a critical role in sensory gating and sensory inhibition (Buzsáki & Draguhn, 2004; Foxe & Snyder, 2011). Specifically, the bottom-up posterior→frontal propagation of alpha activity has been found to regulate large-scale networks such that unwanted information (e.g., sensory distracters) can be blocked (Johnson, et al., 2017), while the suppression of alpha activity would lift the inhibition, permitting sensory inputs to enter conscious awareness as explicit percepts (Emrich, Riggall, Larocque, & Postle, 2013; Postle, 2016; Sadaghiani & Kleinschmidt, 2016; Sadaghiani, et al., 2012). Here, we observed that attenuated posterior→frontal alpha connectivity correlated with both intrusive re-experiencing and trauma memory (for both combat and non-combat odors) in the combat veterans. Furthermore, as indicated by our mediation analysis, attenuated posterior→frontal alpha connectivity contributed to intrusive re-experiencing via heightened olfactory trauma memory. To note, we also tested alternative mediation models, with alpha connectivity or intrusive re-experiencing as the mediator of the three-way correlations, which confirmed this specific mediation by trauma memory (see Supplemental Materials).

This attenuation of posterior→frontal alpha connectivity was observed during the resting state, highlighting its intrinsic nature and thereby close link to spontaneous memory recall. Furthermore, this attenuated alpha connectivity was not associated with emotional response to odors, accentuating its specific link to sensory-based trauma memory activation. As such, not only trauma-related but also mundane cues could break into consciousness as vivid sensory percepts and imagery, driving intrusive symptoms. In addition, as mundane cues are purportedly non-threatening and thus unlikely to activate the “alarm” defense system, they may not elicit sufficient top-down executive control. Therefore, unchecked by attention or voluntary inhibition, non-trauma cues (e.g. non-combat odors) may be uniquely positioned to “sneak” into downstream processes of memory and consciousness, evoking intrusive re-experiencing. From a clinical perspective, such mundane triggers for intrusion would account for the so-called “ease of triggering” and pervasive, uncontrollable nature of intrusive symptoms in PTSD (Ehlers, 2010).

That this alpha oscillation effect emerged in the long-range alpha connectivity as opposed to local alpha power emphasizes the relevance of sensory→frontal inhibitory regulation, relative to focal modulation of sensory cortical excitability, in trauma memory activation. The posterior→frontal propagation of alpha oscillations is known to originate in the occipital (and parietal) visual cortex to target various frontal regions, including the anterior insula and anterior cingulate cortex (Hillebrand, et al., 2016; Johnson, et al., 2017; Sadaghiani, et al., 2010; Tang, et al., 2007). The anterior insula and anterior cingulate cortex are postulated to be key neural substrates of the sensory-based system of trauma memory (Brewin, 2014) in addition to central components of the salience network critical for salience detection and vigilant response driven by bottom-up sensory inputs (Menon, Woodward, Pomarol-Clotet, McKenna, & McCarthy, 2005). Akin to the inhibitory function of alpha oscillations, attenuated alpha activity is coupled with heightened salience network activity (Ros, et al., 2013; Sadaghiani, et al., 2010), which are found to coalesce in PTSD (Clancy, et al., 2017; Lanius, Frewen, Tursich, Jetly, & McKinnon, 2015; Sripada, et al., 2012). Together, this conjunction of neural substrates suggests that the combination of weakened alpha posterior→frontal connectivity and heightened salience network activity in PTSD would precipitate pathological overactivation of sensory-based trauma memory, resulting in vivid, sensory-rich intrusions of traumatic events.

While intrusive re-experiencing takes place in all sensory modalities, the visual system has been most extensively studied (Brewin, et al., 2010). Our use of odors as sensory cues allowed us to interrogate the relatively understudied olfactory system in relation to this symptom cluster. Human olfaction is deeply intertwined with memory (Wilson & Stevenson, 2003). Moreover, olfactory cues are potent elicitors of involuntary memory, as famously described by Proust (Chu & Downes, 2000; Herz & Engen, 1996). Accordingly, the extant literature indicates that odors are strong triggers of trauma memories in PTSD (Cortese, et al., 2015; Kline & Rausch, 1985; Vermetten & Bremner, 2003; Vermetten, et al., 2007). In addition, the olfactory neuroanatomy is intricately connected with the insula and anterior cingulate cortex (Carmichael, Clugnet, & Price, 1994; Seubert, Freiherr, Djordjevic, & Lundstrom, 2013), which would receive intensified bottom-up input from the primary olfactory cortex in an anxious state (Krusemark, Novak, Gitelman, & Li, 2013). Consistent with these ideas, our finding of olfactory trauma memory mediating the association between sensory disinhibition and intrusive trauma re-experiencing implies that fleeting whiffs of odorous air in the environment may inadvertently trigger strong trauma memories and intrusive symptoms. As such, increased attention to this sensory modality is warranted not only in basic research but also in clinical application such as the incorporation of odor-based exposure interventions.

Negative intrusions are observed in multiple psychiatric disorders. However, the strong perceptual vividness and rich, concrete sensory features of intrusive re-experiencing in PTSD sets these symptoms apart from abstract intrusive thoughts that are common in depression and other anxiety disorders (Brewin, et al., 2010). This sensory-rich quality would render the intrusions highly experiential and, to a great extent, perceived as occurring in the present. This distinct nature of PTSD-related intrusions strongly implicates sensory aberrations and promotes sensory-based conceptualization of this disorder. Towards that end, our findings provide a mechanistic, sensory account of PTSD pathology: deficits in bottom-up sensory inhibition allow for mundane sensory inputs from the environment to spontaneously activate sensory representations of trauma memories, evoking involuntary recall and re-experiencing of traumatic events.

## Acknowledgement

This research was supported by the National Institute of Mental Health grants R01MH093413 (W.L.), the FSU Chemical Senses Training (CTP) Grant Award T32DC000044 (K.C.) from the National Institutes of Health (NIH/NIDCD), and the Subaward of U.S. Army award W81XWH-10-2-018 (N.S.), which does not necessarily represent the views of the Department of Defense, Department of Veterans Affairs, or the United States Government, nor does it constitute or imply endorsement, sponsorship, or favoring of the study design, analysis, or recommendations.

## SUPPLEMENTAL INFORMATION

### SUPPLEMENTAL RESULTS

#### No association between alpha power and olfactory memory or emotional response

An omnibus rANCOVA (Category x Response x alpha power) of olfactory response revealed no main effect of alpha power on ratings (*p* = .359) or interactions with Category or Response (*p*’s > .280).

#### Trauma memory for odors associated with intrusive re-experiencing symptoms over and beyond emotional response

A multiple regression was performed entering trauma memory for and emotional response to odors (combat and non-combat) as regressors on intrusive re-experiencing symptoms. The overall model was significant (*F*_2, 34_ = 5.47, *p* = .007), and there was a unique association between intrusive re-experiencing and trauma memory (*sr* = .45, *p* = .007) over and beyond emotional response (*sr* = .12, *p* = .506).

Another multiple regression was performed entering trauma memory for combat and non-combat odors as separate regressors on intrusive re-experiencing symptoms to assess their unique roles in intrusive re-experiencing symptoms. The overall model was significant (*F*_2, 34_ = 6.20, *p* = .005), with a unique association between intrusive re-experiencing and trauma memory for non-combat odors (*sr* = .36, *p* = .036) over and beyond combat odors (*sr* = .05, *p* = .776).

#### Frontal→posterior alpha connectivity was not related to intrusive re-experiencing symptoms

To demonstrate the specificity of the posterior→frontal direction of alpha connectivity on intrusive re-experiencing symptoms, multiple regressions were performed using frontal → posterior alpha connectivity. Neither left (*sr* = −0.07, *p* = .537) nor right (*sr* = −0.12, *p* = .264) were associated with intrusive re-experiencing symptoms. Supplemental Figure 1 further demonstrates the dominance of the posterior→frontal direction of alpha connectivity at rest, relative to frontal→posterior, substantiating its integral role in regulating resting state networks responsible for the orchestration of sensory and cognitive processes.

#### Alternative Mediation Models

Alternative mediation models were performed to evaluate the specificity of the relationship between alpha connectivity, trauma memory, and intrusive re-experiencing symptoms. Specifically, attenuated alpha connectivity was not related to greater trauma memory through more severe intrusive re-experiencing symptoms (indirect effect: *β* = 1.32, SE = 1.32, 95% CI = [−0.362, 5.309]). Similarly, more intrusive re-experiencing symptoms were not related to greater trauma memory through attenuated alpha connectivity (indirect effect: *β* = 1.32, SE = 1.44, 95% CI = [−.483, 5.558]). Other alternative iterations of the mediation model did not yield statistical significance, highlighting the specificity of the relationship of alpha connectivity on intrusive re-experiencing symptoms mediated by biased trauma memory of sensory cues.

